## Supplemental figures for "Human visual gamma for color stimuli: When LGN drive is equalized, red is not special"

### Figure Supplement

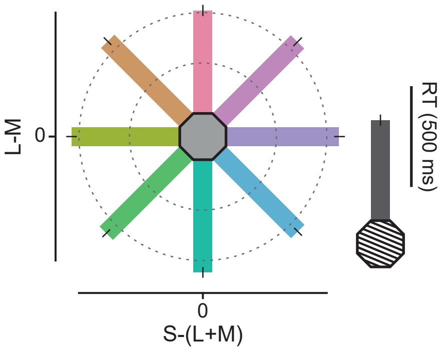


Figure 2-figure supplement 1. Reaction time results: Average reaction times for all stimuli. Error bars represent 95% CIs over participants, bar orientation represents stimulus orientation in DKL space. The length of the scale bar corresponds to the distance from the edge of the hexagon to the outer ring.

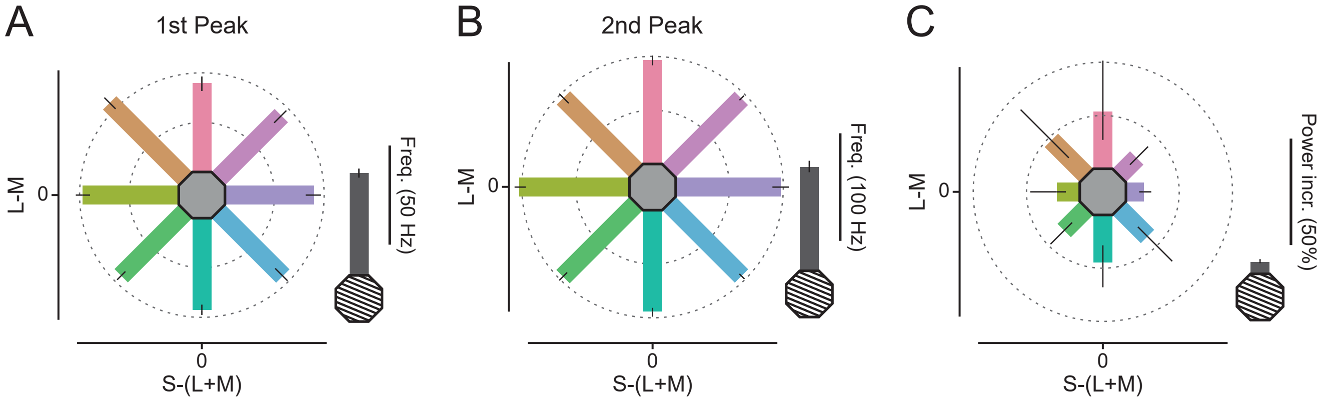


Figure 3-figure supplement 1. Spectral measures: (A) Average induced gamma-peak frequency of the lower gamma peak for all stimuli. Bar orientation represents stimulus orientation in DKL space. In grey, the same is shown for the grating stimulus. (B) Same as A, but for the upper gamma peak frequency. (C) Same as A, but for stimulus-induced power at the upper gamma peak frequency. In all panels, error bars represent 95% CIs over participants and the length of the scale bar corresponds to the distance from the edge of the hexagon to the outer ring.

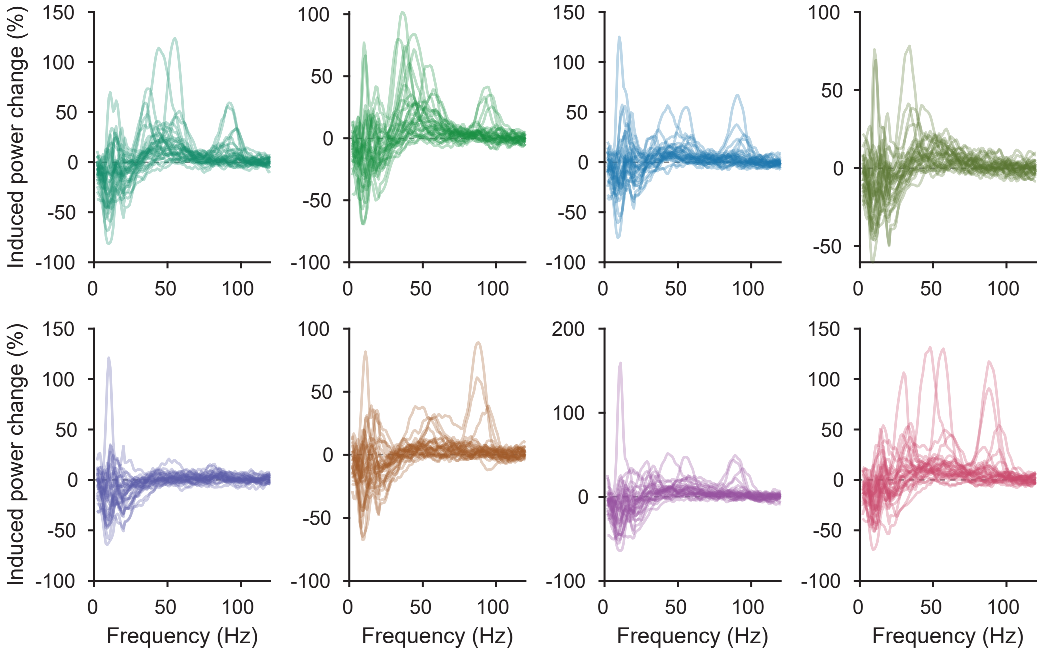


Figure 3-figure supplement 2. Individual participant spectra: Stimulus-induced power changes over baseline (averaged over V1 dipoles) for all 30 participants and the eight presented colors.
